# Role and patterns of butterflies and hawkmoths in plant-pollinator networks at different elevations and seasons in tropical rainforests of Mount Cameroon

**DOI:** 10.1101/2021.02.16.431477

**Authors:** Jan E.J. Mertens, Lucas Brisson, Štěpán Janeček, Yannick Klomberg, Vincent Maicher, Szabolcs Sáfián, Sylvain Delabye, Pavel Potocký, Ishmeal N. Kobe, Tomasz Pyrcz, Robert Tropek

## Abstract

1. Butterflies and moths are well-visible flower visitors. Nevertheless, almost no quantification of their role in plant-pollinator interactions exists at a community level, especially from tropical rainforests. Moreover, we have virtually no knowledge on environmental and other factors affecting lepidopteran flower visits.
2. We focused on the role of butterflies and hawkmoths as flower visitors in tropical rainforests of Mount Cameroon, especially on its elevational and seasonal changes. We also analysed their preferences to selected floral traits, with a specific focus on pollination syndromes.
3. We video-recorded flower visitors of 1,115 specimens of 212 plant species (>26,000 recording hrs) along the complete elevational gradient of rainforests in two main seasons, and compared frequencies of flower-visiting lepidopterans to other visitors. We compared characteristics of plant-lepidopteran networks among elevations and seasons, and analysed patterns of selected lepidopteran traits. Finally, we analysed inter-family differences in their floral preferences.
4. Altogether, we recorded 734 flower visits by 80 butterflies and 27 hawkmoth species, representing only ~4% of all 18,439 flower visits. Although lepidopterans visited only a third species, they appeared key visitors of several plants. The most flower visits by lepidopterans were recorded in mid-elevations and dry season, mirroring the general patterns of lepidopteran diversity. The networks showed no apparent elevational or seasonal patterns, probably because of the surprisingly high specialisation of interactions in all networks. Significant non-linear changes of proboscis and forewing lengths were found along elevation, and long-proboscid hesperiid butterflies visited flowers with longer tubes or spurs. Substantial differences in floral preferences were found between sphingids, and papilionid, nymphalid and lycaenid butterflies, revealing importance of nectar production, floral size and shape for sphingids, and floral colour for butterflies.
5. Butterflies and hawkmoths were confirmed as relatively minor visitors of tropical forest flowers, although they seemed crucial for pollination of some plant species. Moreover, the revealed floral preferences and trait-matchings confirmed a potential of some lepidopteran families to drive floral evolution in tropical ecosystems.

## Introduction

Recently, pollination research shifted from detailed studies of single pollination systems to network approaches. Nevertheless, most complex studies of individual pollinator groups’ role in plant-pollinator networks have focused on bees or hoverflies (e.g. Classen et al., 2020; Klecka et al., 2018), whilst the other flower visitors have often been excluded or side-lined. Although some less abundant groups play important roles in pollination systems, as secondary pollinators, nectar thieves and competitors, or even as key pollinators of specialised plants (Hahn & Brühl, 2016; Martínez-Adriano et al., 2018; Mertens et al., 2020; Ollerton, 2017; Wardhaugh, 2015), their importance in plant-pollinator networks remains understudied, especially in tropical forests.

Compared to bees and flies, butterflies and hawkmoths represent minor pollinators in probably all terrestrial ecosystems (Ollerton, 2017; Wardhaugh, 2015). Both groups are often regarded as generalised nectar feeders visiting all available nectar-rich flowers (Johnson et al., 2017; Willmer, 2011). Even hawkmoths, considered as efficient pollinators strongly affecting floral evolution already since Darwin (1862), were recently revealed as opportunistic nectar thieves of many flowers (e.g. Fox et al., 2015; Martins & Johnson, 2013). However, some butterflies (e.g. Arroyo et al., 2007; Mertens et al., 2020; Santos et al., 2020) and moths (e.g. Fleming & Holland, 1998; Hahn & Brühl, 2016; Johnson et al., 2017; Skogen et al., 2019) are key pollinators of specialised plants.

Individual lepidopteran groups differ in their morphological and behavioural adaptations to pollination. Among butterflies, papilionids, pierids, and some groups of nymphalids and hesperiids use their long proboscis to feed on nectar from deep flowers, whilst many lycaenids, riodinids, and some smaller clades within the mentioned families bear small proboscis unable to reach nectar in specialised flowers (Corbet, 2000; Tiple et al., 2009). In moths, besides highly specialised long-proboscid groups, such as most sphingids and noctuids, adults of many groups have dysfunctional or even no proboscis (Willmer, 2011). Such differences hamper any attempts at quantifying the general pollination role of lepidopterans. Especially at the community level, the relative importance of butterflies and hawkmoths as pollinators is understudied.

Plants also differ in their adaptation to butterfly or moth pollination. The pollination syndrome hypothesis (Faegri & van der Pijl, 1979) expects some plants to evolve certain traits to attract the two groups. Psychophily hypothesises the adaptation for butterfly-pollination, whilst sphingophily defines hawkmoth-pollinated flowers and is distinguished from phalaenophily, i.e. pollination by any other moths (Faegri & van der Pijl, 1979; Willmer, 2011). Consistently, butterflies and hawkmoths should prefer large conspicuous flowers or inflorescences (Arroyo et al., 2007; Mitchell et al., 2015). Nocturnal hawkmoths rely equally on colour and scent when foraging, often preferring light colours (such as white or cream) better distinguishable in dark, and strong sweet scents (Glover, 2011; Kelber et al., 2003). This is in contrast with butterflies typically preferring bright flower colours, such as red or orange, above scent (Ômura & Honda, 2005), although sweet and fruity scents were also included into the psychophily (Willmer, 2011). Nevertheless, the colour preference strongly varies among butterfly families and species (Pohl et al., 2011; Yurtsever et al., 2010). Their size and proboscis length also influence flower preferences (Tiple et al., 2009). Small short-proboscid lycaenids avoid long-tubed flowers but can visit small solitary flowers, long-proboscid papilionids or pierids, often larger and more energy-demanding, prefer massed nectar-rich flowers (Corbet, 2000; Tiple et al., 2009). Long-proboscid hawkmoths can visit both long and short tubed flowers (Johnson et al., 2017).

Elevation and seasonality, representing various environmental and ecological gradients, influence patterns in biotic interactions (Klomberg et al., 2020; Poisot et al., 2015). The role, relative proportions in communities, and specific adaptations of pollinator groups shift under differing environmental conditions, such as temperature, solar radiation, and precipitation (e.g. Ollerton et al., 2006; Klomberg et al., 2020). Unfortunately, neither elevational nor seasonal patterns of tropical lepidopteran role in pollination networks were studied, except a few case studies of individual plant species (e.g. Mertens et al., 2020). However, we can expect some correlations of their role in networks with their general diversity patterns. We have no community-wide studies on characteristics of these lepidopteran-plant pollination networks in any tropical area.

Our study focuses on flower-visiting butterflies and hawkmoths, the two relatively minor groups of pollinators often overlooked in network studies, yet easily identifiable. The primary Afrotropical rainforests covering Mount Cameroon from nearly sea level to the natural timberline offer a unique elevational gradient, with distinct dry and wet seasons. Based on rich community-wide datasets sampled along the elevational gradient and during the two seasons, we set the following aims: (1) To evaluate the role of flower-visiting butterflies and hawkmoths in plant-pollinator networks and understand how elevation and seasonality affect their relative importance in pollination communities. (2) To analyse potential elevational and seasonal changes in structure of the pollination networks, with a specific focus on specialisation. (3) To assess the butterfly and hawkmoths preferences to floral traits, as well as to test potential trait-matching between flowers and their visitors. (4) To test potential relationship of proboscis length and specialisation of butterflies and moths in visited flowers. We hypothesise that butterflies and hawkmoths represent only a small proportion of the flower-visiting communities, although we expect it will be higher in lowlands and in dry season where and when both groups are more abundant and diverse on Mount Cameroon (Maicher et al., 2018, 2020). We also expect higher specialisation in communities with more flower-visiting lepidopteran species, such as in some other insect-plant interactions (e.g. Rasmann et al., 2018). We expect both groups to be important pollinators of some specialised plants. We hypothesise preferences to some traits previously included in the psychophilous and sphingophilous syndromes, although we expect some predicted traits to be less important. We also expect substantial differences in the mentioned aims and hypotheses among lepidopteran families.

## Materials and Methods

### Study area

Mount Cameroon (4,095 m a.s.l.) is an active volcano in the Southwest Region, Cameroon, West/Central Africa. Primary tropical rainforests cover its southwestern flanks, where the study was performed, from lowlands (above human encroachment at ca. 300 m a.s.l.) up to the natural timberline (ca. 2200 m a.s.l.). As the mountain is located within the ‘Guinean forests of West Africa’ biodiversity hotspot, it holds an extraordinarily high biodiversity of numerous taxa, including butterflies (Larsen, 2005), hawkmoths (Ballesteros-Mejia et al., 2013), and plants (Cheek et al., 1996). The mountain belongs among the wettest places in the world and experiences distinct dry (December–February) and wet seasons (June–September; Maicher et al., 2018, 2020)). The Atlantic Ocean-facing southwestern lowlands receive large amounts of rainfall (>12,000 mm annually), most of which during the wet season (>2,500 mm monthly), and rarely any rain during the dry season (Maicher et al. 2020). To characterise changes in plant-pollinator interactions along elevation and season, we studied four sites on the southwestern slope at 650, 1,100, 1,450 and 2,200 m a.s.l. The butterfly and hawkmoth species richness data includes additional sampling sites at 30, 350 and 1,850 m a.s.l. (Table 1). For more details on the study sites, see Maicher et al. (2020).

**Table 1.** Sites on Mount Cameroon sampled for butterflies and sphingids. ‘n.a.’ states for data not available for particular sites.

| Site |  |  |  | Sampled period |  | Number of all species |  | Species in pollination networks (dry / wet seasons) |  |  |  |
| --- | --- | --- | --- | --- | --- | --- | --- | --- | --- | --- | --- |
| Elevation<br>(a.s.l.) | Latitude | Longitude | Vegetation<br>type | Checklist | Networks<br>(dry/wet) | Butterflies | Sphingids | Butterflies | Sphingids | All plants | Visited plants |
| 30 m | N 03.9818° | E 09.2625° | Coastal forest | Dec 2014, Jan 2015,<br>May 2015,<br>Oct 2017 | n.a. | 282 | 5 | n.a. | n.a. | n.a. | n.a. |
| 350 m | N 04.0899° | E 09.0517° | Mosaic of primary<br>and secondary<br>lowland forest | Dec 2014,<br>Apr 2015,<br>Jan/Feb 2016 | n.a. | 189 | 28 | n.a. | n.a. | n.a. | n.a. |
| 650 m | N 04.1022° | E 09.0630° | Primary lowland<br>forest | Nov/Dec 2014,<br>Apr 2015,<br>Jan/Feb 2016 | Jan 2018 /<br>Aug 2018 | 189 | 20 | 32 / 14 | 5 / 6 | 62 / 42 | 19 / 11 |
| 1,100 m | N 04.1175° | E 09.0709° | Upland forest<br>disturbed by<br>elephants | Dec 2014, Jan 2015,<br>Apr 2015,<br>Jan/Feb 2016 | Feb 2018 /<br>Sep 2018 | 161 | 8 | 38 / 7 | 7 / 4 | 61 / 32 | 25 / 12 |
| 1,450 m | N 04.1443° | E 09.0717° | Submontane forest<br>disturbed by<br>elephants | Nov 2016,<br>Feb 2017,<br>Apr/May 2017 | Feb 2017 /<br>Sep 2017 | 64 | 7 | 13 / 7 | 9 / 4 | 42 / 35 | 17 / 6 |
| 1,850 m | N 04.1453° | E 09.0870° | Montane forest<br>disturbed by<br>elephants | Nov 2016,<br>Feb 2017,<br>Apr 2017 | n.a. | 12 | 7 | n.a. | n.a. | n.a. | n.a. |
| 2,200 m | N 04.1428° | E 09.1225° | Montane forest<br>close to timberline | Nov 2016,<br>Jan/Feb 2017,<br>Apr 2017 | Jan 2017 /<br>Aug 2017 | 13 | 3 | 3 / 0 | 2 / 1 | 22 / 28 | 6 / 2 |
| TOTAL |  |  |  |  |  | 431 | 40 | 80<br>(69 / 25) | 26<br>(19 / 12) | 212<br>(144 / 106) | 71<br>(54 / 26) |

### Study groups and their biodiversity patterns

This study focused on butterflies (Lepidoptera: Papilionoidea) and hawkmoths (Lepidoptera: Sphingidae; hereafter referred to as sphingids). For part of the analyses, butterflies were split up in their families (Hesperiidae, Papilionidae, Pieridae, Lycaenidae, and Nymphalidae; hereafter referred to as hesperiids, papilionids, pierids, lycaenids, and nymphalids). All butterflies and sphingids were represented in the *flower visitation* and *floral preferences* parts of the study. However, the effects of elevation and season on visitor size and proboscis length were analysed for papilionids, hesperiids and sphingids only, because the other groups’ traits were not measured in the field. We actively inventoried Lepidoptera along the complete elevation of Mt. Cameroon (i.e. at seven elevations from 30 to 2,200 m a.s.l.), using the checklist approach. For this purposes, we applied intensive hand-catching (our unpublished data) and bait-trapping (data from Maicher et al. 2020) of butterflies, whereas standardised light-attraction of sphingids (data from Maicher et al. 2020). These data were further supplemented by a few additional species found only in the video recordings described below.

### Flower visitation

We recorded flower-visiting lepidopterans at the four elevations (650, 1,150, 1,450 and 2,250 m a.s.l.), along six transects (200×10 m) per elevation established to characterise the local vegetation heterogeneity (Klomberg et al., 2020). Along those transects, we recorded flower visitors of all plant species flowering during our fieldwork (two weeks during dry and wet seasons at each elevation; Table 1) using security cameras with IR night-vision (Vivotek IB8367RT). We positioned the cameras 0.5–1.5 m from the flowers or inflorescences and camouflaged their surfaces. We recorded flowering plants at all vegetation layers from understorey to canopy, using ladders and tree climbing to reach higher strata. Five individuals of each plant species were recorded, each for a 24hour session. The individual replicates were separated in space (different transects) and time (different days). During the first week, we added any plant species that have been or just started flowering. The second week served towards completing the necessary five replicates and no more species were added to the study. Whenever insufficient individuals flowered along the transects, we searched the adjoining area.

We observed all flower visitors from the video recordings either through semi-autonomic motion detection with Motion Meerkat 2.0.5 (Weinstein, 2015) when conditions allowed, or manually through sped-up playback. We identified all butterflies and sphingids to (morpho)species using various available literature and our reference collection. For each visiting Lepidoptera, we determined whether they touched the plant’s reproductive organs (anthers, stigmata, or both) to distinguish potential pollinators from other visitors.

The recorded interactions were used to reconstruct interaction networks among flowering plants and visiting lepidopterans (hereafter simplified to *plant-lepidopteran networks* or *networks*) for each elevation and season (i.e. eight networks). We used visitation frequency (i.e. number of interactions of each species per plant species during 24h) in each of the eight networks. This controls for differences in total recording time between plant species in the few cases we failed to find enough replicates or the recordings were shorter because of dying flowers or technical failures.

To visualise and characterise the eight plant-lepidopteran networks, we used the *bipartite* package (Dormann et al., 2009) in R 3.5.3 (R Core Team, 2019). We quantified network *connectance* (Jordano, 1987), network-level *H2’ specialisation* (Blüthgen et al., 2006), *Q modularity* (Dormann & Strauss, 2014), and *NODF nestedness* (Almeida-Neto et al., 2008). We calculated each metric firstly including all floral visitors, and secondly only with the subset of visitors touching the plant’s reproductive organs. Because of the highly limited number of replicates (each combination of elevation and season was characterised by a single network), any possible elevational and seasonal patterns of the network characteristics were checked by a direct comparison of values, i.e. without any statistics. Finally, we calculated *d’ specialisation* (Blüthgen et al., 2006) of each lepidopteran species in each network. The relationship of the species-level specialisation of lepidopterans to elevation and season was analysed by a linear mixed-effect model (LMM) with specialisation of individual lepidopteran species in particular networks as a continuous response variable, and with elevation and season as categorical explanatory variables. Individual lepidopteran families were included as a categorical random-effect factor to correct for the inter-family variability.

### Relationship between floral and lepidopteran traits

We measured six floral traits of 174 plant species included in the plant-pollinator networks: *symmetry* (actino- or zygomorphic), prevailing *floral colour*, *corolla width*, floral *tube length* (distance from the flower opening to its base, or tip of the spur when present), and *nectar sugar* (total mass of sugars produced by a flower during 24h; the sampling protocol followed Bartoš et al. 2020).

We measured eight morphometric traits of 1,665 specimens of 130 lepidopteran species (75 hesperiids, 15 papilionids, and 40 sphingids) collected during the project. Directly in the field, we *weighted* fresh specimens and cut their proboscides for later measurement. The collected specimens were mounted and photographed at the Nature Education Centre, Jagiellonian University, Krakow. On these photographs, we measured *forewing length* and *width*, *body length*, and *thorax width*, *lengths of fore*-, *mid*- and *hindleg*, and *proboscis* (Fig. S1) in ImageJ2 (Rueden et al., 2017). We assessed the lepidopteran trait collinearity by multiple regression and selected the proboscis and forewing lengths as the other traits proxies (Table S2).

We analysed patterns of the proboscis and forewing lengths in communities of all lepidopterans measured at different elevations by LMM. The average trait values per species were used as response variable (log-transformed as the data showed a lognormal distribution), elevation as categorical fixed-effect variable, and lepidopteran families as random-effect variable to correct for inter-family variability. Consequently, we analysed elevational and seasonal (fixed-effect variables) differences in the proboscis or forewing lengths (response variables) in flower-visiting lepidopteran species only. This dataset involved 34 measured lepidopteran species recorded during flower visits (19 hesperiids, 7 papilionids, 8 sphingids). In both analyses, we applied AICc (AIC corrected for small samples, Hurvich and Tsai 1993) to select the most plausible models. Due to the high variability in sample sizes, no post-hoc tests were performed. Finally, we tested correlation between the proboscis and forewing length of lepidopterans and their d’ specialisation using Spearman’s rank coefficients.

### Floral preferences

We assessed how the floral preferences to particular floral traits differ among the six focal lepidopteran families by ordination analyses in Canoco 5 (ter Braak & Šmilauer, 2012). All five measured flower traits served as explanatory variables towards the visitation frequencies by lepidopteran species (response variable). Based on the gradient lengths, two RDA models were selected and tested using 999 Monte Carlo permutations (Šmilauer & Lepš, 2014). Firstly, to assess lepidopteran preferences within the whole local community of flowering plants, we included all plant species for which we measured the traits (n=173). Subsequently, we analysed lepidopteran preferences only among the visited plant traits (n=63). Finally, we tested whether lepidopterans with longer proboscides visit flowers with longer corolla tubes by correlating the average proboscis length of the 34 measured lepidopteran visitors with the corolla tube length of the visited plant species using Spearman’s rank coefficient.

## Results

Altogether, we recorded 431 butterfly and 40 sphingid species on Mount Cameroon (Table 1; Fig. 1a). Nymphalids comprised the most species across the gradient, followed by lycaenids in the lowest elevations and hesperiids at the mid-elevations (650 m to 1,450 m a.s.l.). The total species richness of lepidopterans, as well as of lycaenids, showed gradual decrease along elevation (Fig 1a), all other butterflies showed the low plateau pattern (*sensu* McCain and Grytnes 2010), whilst sphingids peaked at 350 m a.s.l. (Fig. 1a).

**Figure 1.**
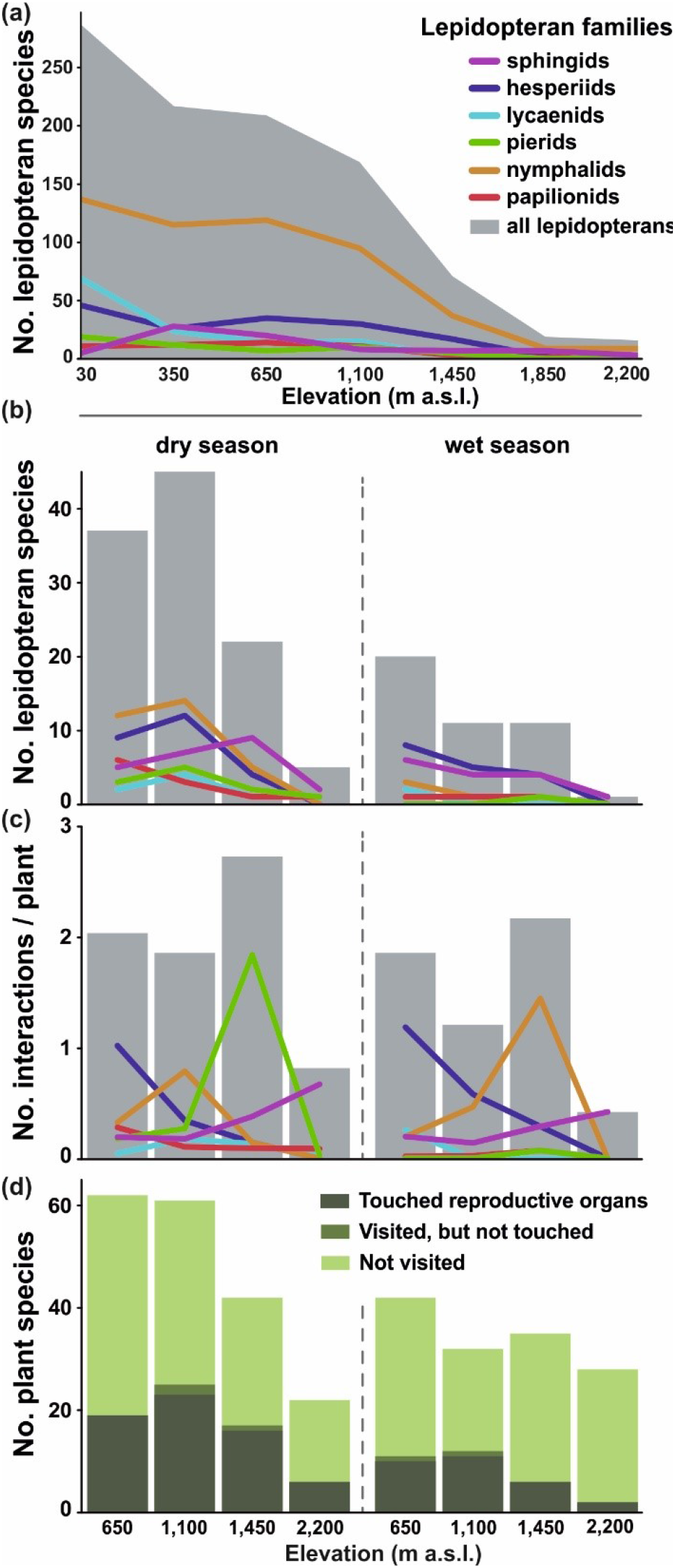
Overview of (a) lepidopteran species richness along the elevational gradient of Mount Cameroon, (b) total number of flower-visiting lepidopteran species at each elevation and season, interaction frequency per plant and 24hrs, and plant species where lepidopterans touched reproductive organs, where lepidopterans visited without touching any reproductive organs, and where lepidopteran visits were not recorded. Grey shading in (a-c) denotes the sum of all lepidopteran taxa, coloured lines represent particular families.

Altogether, we recorded 1,115 individuals of 212 flowering plant species for a combined 26,138 hours (~2.98 years) of video footage, during which we observed 734 individuals of 80 butterfly and 27 sphingid species visiting 71 plant species. These visits represented 4% of all 18,439 flower visits recorded on the observed plants. Bees, flies, beetles, and other moths were more common flower visitors than butterflies (Klomberg et al., 2020). Wasps, nectarivorous birds and carpenter bees were more common visitors than sphingids, followed by cockroaches and mammals (Klomberg et al., 2020). Still, butterflies and sphingids were among the two most common flower visitors for some plant species, such as *Scadoxus cinnabarinus* (Amaryllidaceae), *Distephanus biafrae* and *Melanthera scandens* (both Asteraceae), and *Cordia aurantiaca* (Boraginaceae) for butterflies; and *Anthocleista scandens* (Gentianaceae) for sphingids (Table S4). From these, 700 lepidopteran visitors touched the plant reproductive organs (see Table S3 for a taxonomic and spatiotemporal overview). Due to the small difference between ‘pollinators’ and ‘all visitors’, results of our analyses with both datasets were nearly identical, and we only report all visiting records, i.e. the visitors’ point of view.

We recorded the highest species richness of both lepidopteran flower visitors and lepidopteran-visited plants at 1,100 m a.s.l. during dry season (Table 1; Fig. 1b). Species richness of visiting lepidopterans decreased towards the higher elevations and during wet season; at 2,200 m a.s.l. during wet season we recorded only a sphingid species visiting two plant species (Fig. 1b). In accordance, species richness of all flowering plants decreased towards the higher elevations and in wet season. Yet, the highest elevation was species-richer during wet than during dry season. Overall, lepidopterans visited a lower proportion of flowering plant species during the wet season (wet season mean = 0.204 (±0.114) vs. dry season mean = 0.334 (±0.054); Fig. 1d).

The visitation frequency varied among the visitor families (Fig. 1c). Hesperiids were less frequent towards the higher elevations in both seasons. Papilionids followed such pattern in dry season but represented a generally small proportion of visitors in wet season. Lycaenids were generally uncommon flower visitors with only small spatiotemporal differences. Pierids expressed a peak in frequency at 1,450 m a.s.l. during dry season (driven by *Mylothris* cf. *hilara* frequently visiting *Distephanus biafrae*). Nymphalids expressed a similar peak at 1,450 m a.s.l. during wet season (driven by *Vanessula milca* visiting *Melanthera scandens*). Finally, sphingids visited flowers more frequently towards the higher elevations in both seasons (Fig. 1c).

We found no apparent general pattern in turnover of flower-visiting lepidopterans and lepidopteran-visited plants among the studied elevations and seasons (Fig. S2). The higher elevations shared less plant species with the lower elevations as well as between each other. The visitor community shared most species between 1,100 m and 1,450 m, followed by 1,450 m and 2,200 m a.s.l. (Fig. S2).

The plant-lepidopteran networks decreased in size towards the higher elevations and wet season, although the generally largest network was recorded at 1,100 m a.s.l. in dry season (Fig. 2). The trends in the network characteristics were minor or none, except NODF nestedness. Network connectance slightly increased along the elevational gradient and remained similar between seasons. Q modularity slightly decreased towards the higher elevations and during wet season. Whilst NODF nestedness increased along elevation during dry season and showed an opposite trend during wet season. H2’ specialisation slightly increased along elevation during wet season, whilst no pattern was observed during dry season (Fig. 3a-e).

**Figure 2.**
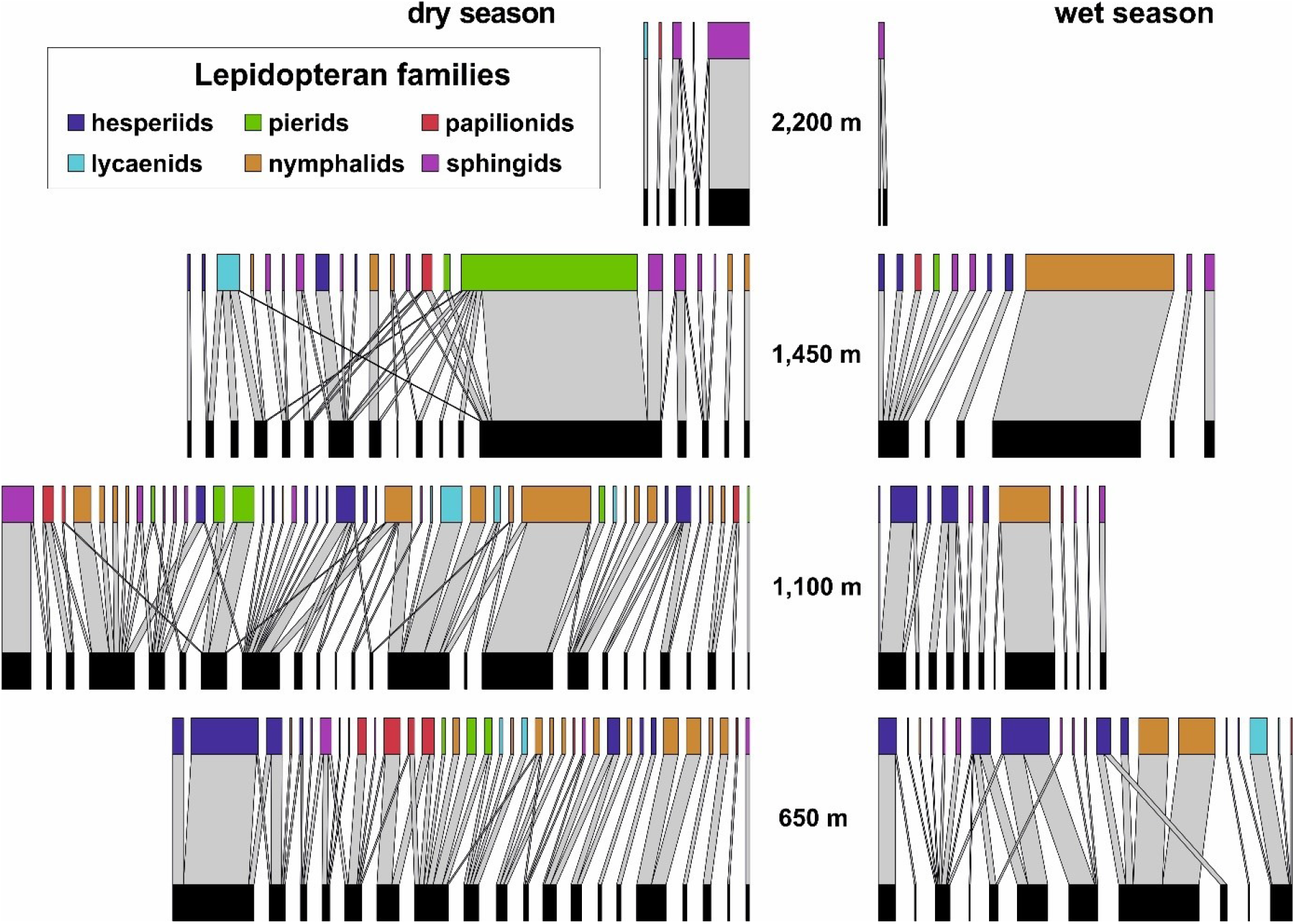
Bipartite networks of plant-lepidopteran interactions along the elevational gradient of Mount Cameroon. The upper nodes visualise flower-visiting lepidopteran species, distinguished by colour for families, whilst the lower nodes represent lepidopteran-visited plant species. The total width of each network approximates their relative size corrected for the sampling effort (visitation frequency per 24hrs of video-recording). The width of individual links (light grey) represents the relative frequency of interactions between visiting lepidopterans and visited plants within each network.

**Figure 3.**
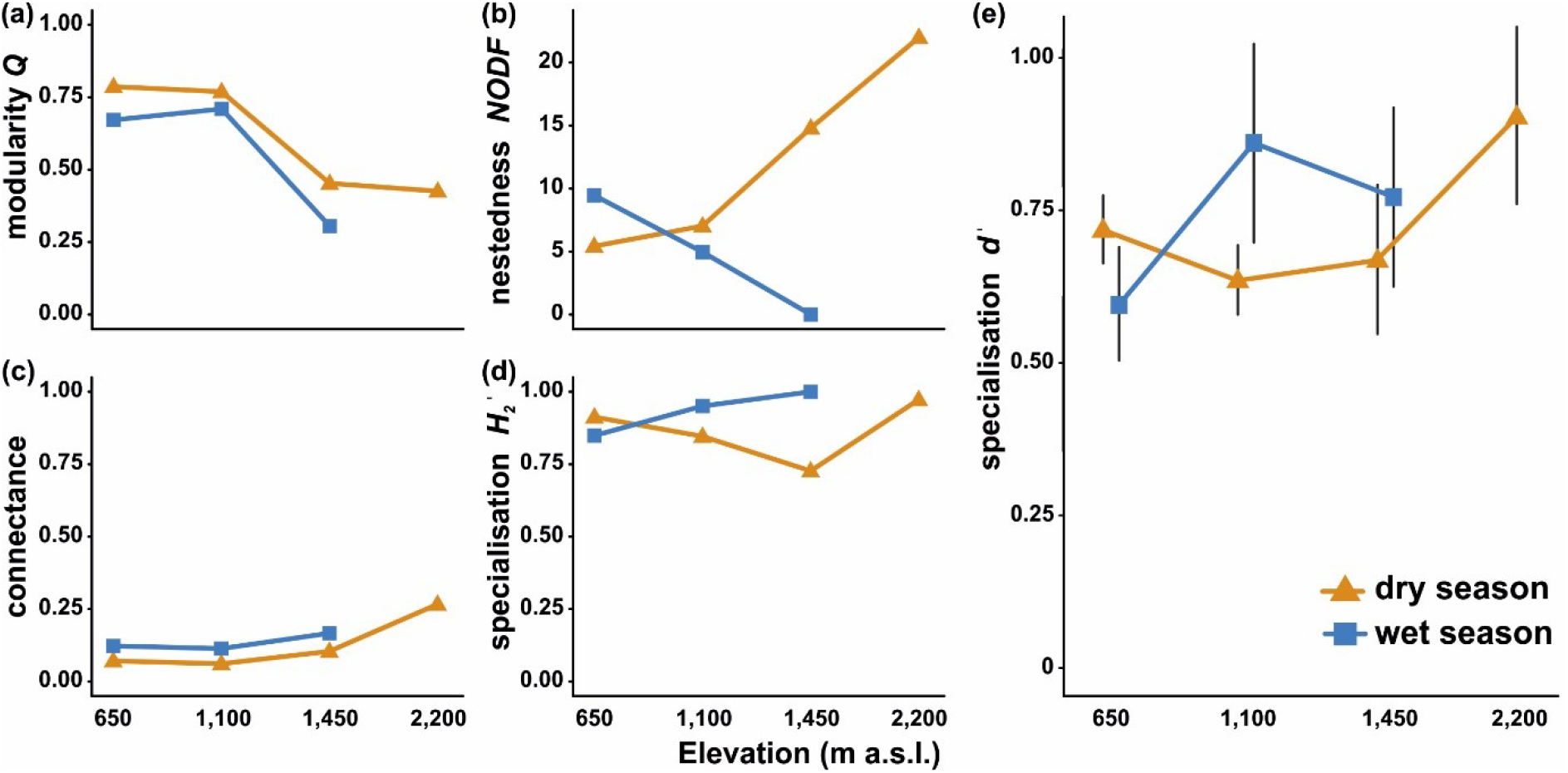
Metrics of plant-lepidopteran networks on Mount Cameroon, comparatively for each elevation and season. The symbols depict arithmetic means in all plots, whilst error bars in (e) represent 95% confidence intervals.

The studied lepidopteran families did not significantly differ in *d’* specialisation (*F* = 0.865, *p* = 0.508), nor there was any significant difference in *d’* specialisation between sphingids and all butterflies (*F* = 0.287, *p* = 0.593). Model comparisons of the effects of elevation, season, and their interaction showed that the interaction effect of both factors is the most plausible descriptor of the observed patterns in *d’* specialisation (Fig. 3e; Table 2).

**Table 2.** Comparison of the effects of season, elevation, and their interaction on *d’* specialisation of flower-visiting lepidopterans on Mount Cameroon. LMM with the lepidopteran families as random-effect variable were applied; models with *ΔAICc* ≤ 2 were considered comparable.

| **All lepidopterans** | *residual*<br>*df* | *residual*<br>*deviance* | $\Delta AICc$ | *weight* | $R^2_{adj}$ |
| --- | --- | --- | --- | --- | --- |
| season | 149 | 7.00 | 11.1 | 0 | 0 |
| elevation | 147 | 6.77 | 10.3 | 0.01 | 0.017 |
| **season × elevation** | **144** | **6.05** | **0** | **0.99** | **0.102** |

### Patterns in lepidopteran morphological traits

Both proboscis and forewing lengths differed significantly among the families (proboscis length: *χ*^*2*^ = 12.15, *df* = 2, *p* = 0.002; forewing length: *χ*^*2*^ = 31.95, *df* = 2, *p* < 0.001). Sphingids had on average the longest proboscides, followed by papilionids and hesperiids. Papilionids had on average the longest forewings, followed by sphingids and hesperiids (Table S5). None of the three families showed any significant patterns in their proboscis or forewing lengths along the elevational gradient (Table S6). However, when analysing only the flower-visiting species, elevation became the most plausible descriptor of the U-shaped patterns in their proboscis and forewing lengths (Table 3; Fig. 4a,b). Even though only three species were measured at the highest elevation, omitting them had no substantial effect on the model strength (Table S7). The proboscis or forewing lengths had no significant effect on *d’* specialisation (proboscis length: *p* = 0.38, *R*^*2*^ = 0.16; forewing length: *p* = 0.63, *R*^*2*^ = 0.086).

**Table 3.** Linear model comparison of the individual effects of season and elevation, and their interaction, on proboscis and forewing length of lepidopterans on Mount Cameroon. ‘*res. df*’ and ‘*res. dev.*’ represent the residual’s degrees of freedom and deviance, respectively.

| Proboscis length | *res. df* | *res. dev.* | $\Delta AICc$ | *weight* | $R^2_{adj}$ |
| --- | --- | --- | --- | --- | --- |
| season | 51 | 73.069 | 3.2 | 0.16 | 0.007 |
| **elevation** | **49** | **64.940** | **0** | **0.78** | **0.114** |
| season × elevation | 45 | 58.864 | 5.2 | 0.06 | 0.180 |
| Forewing length | *res. df* | *res. dev.* | $\Delta AICc$ | *weight* | $R^2_{adj}$ |
| season | 52 | 15.631 | 9.4 | 0.01 | 0 |
| **elevation** | **50** | **1.276** | **0** | **0.88** | **0.062** |
| season × elevation | 46 | -5.802 | 4.1 | 0.11 | 0.090 |

**Figure 4.**
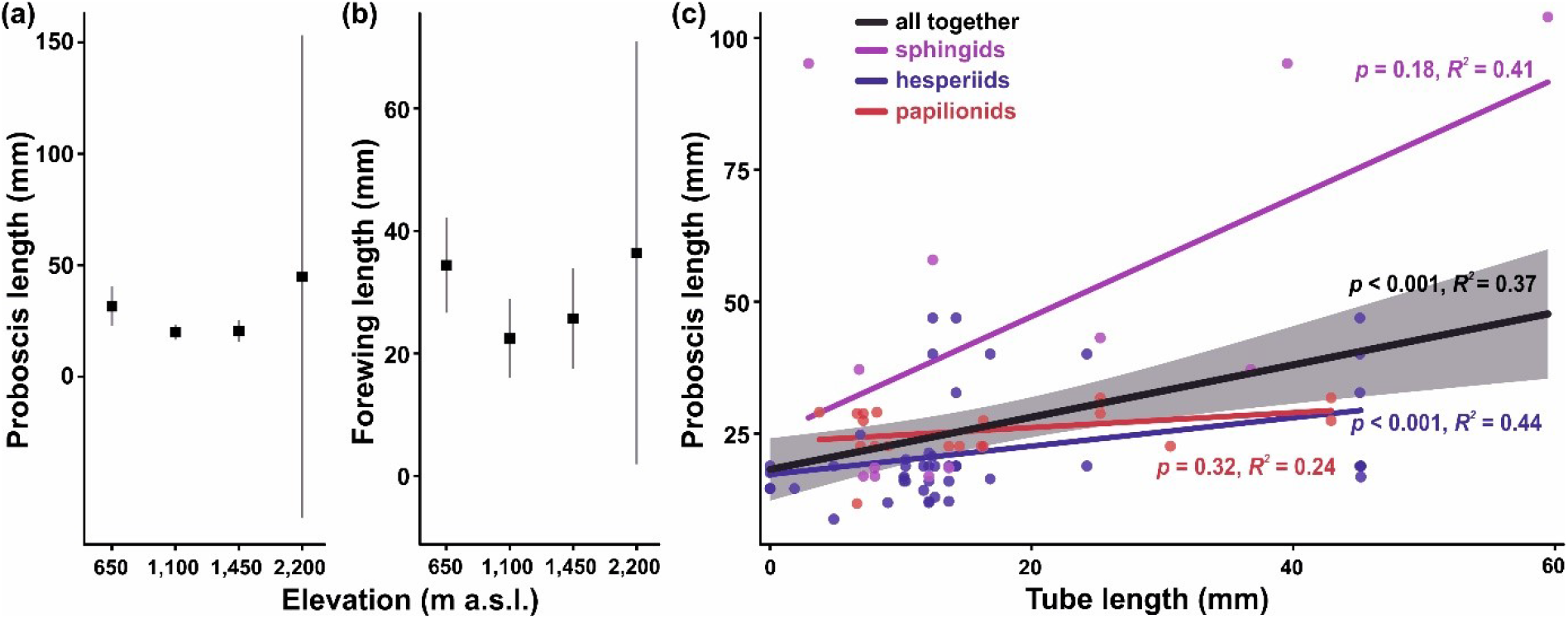
(a) Proboscis and (b) forewing lengths of flower-visiting lepidopterans on Mount Cameroon. Mean values and 95% confidence intervals are visualised. (c) Spearman correlations of lepidopteran proboscis length and corolla tube length of lepidopteran-visited plants. Each data point represents an interaction between a plant species and a lepidopteran species. The black line visualises correlation of all data (with grey shaded confidence intervals), whilst the coloured lines visualise correlations of individual lepidopteran families.

### Lepidopteran preferences to floral traits

Flower-visiting lepidopterans showed significant preferences towards certain floral traits (Fig. 5). Within the complete flowering plant dataset, the selected floral traits explained 25% variability in the visitation frequency (Fig. 5a). The focal families formed three relatively distinct groups. Sphingids preferred sugar-rich, larger and deeper flowers of purple colour. Papilionids, lycaenids and nymphalids preferred orange flowers, whilst hesperiids and pierids did not express any apparent preferences to floral traits. These preferences were mostly consistent with the analysis including only the visited flowers (Fig. 5b), although hesperiids preferred pink actinomorphic flowers. We also found a significantly positive correlation between lepidopteran proboscis length and corolla tube length of lepidopteran-visited flowers (Fig. 4c). However, from the three lepidopteran families, the observed relationship was only significant for hesperiids when analysed separately (Fig. 4c).

**Figure 5.**
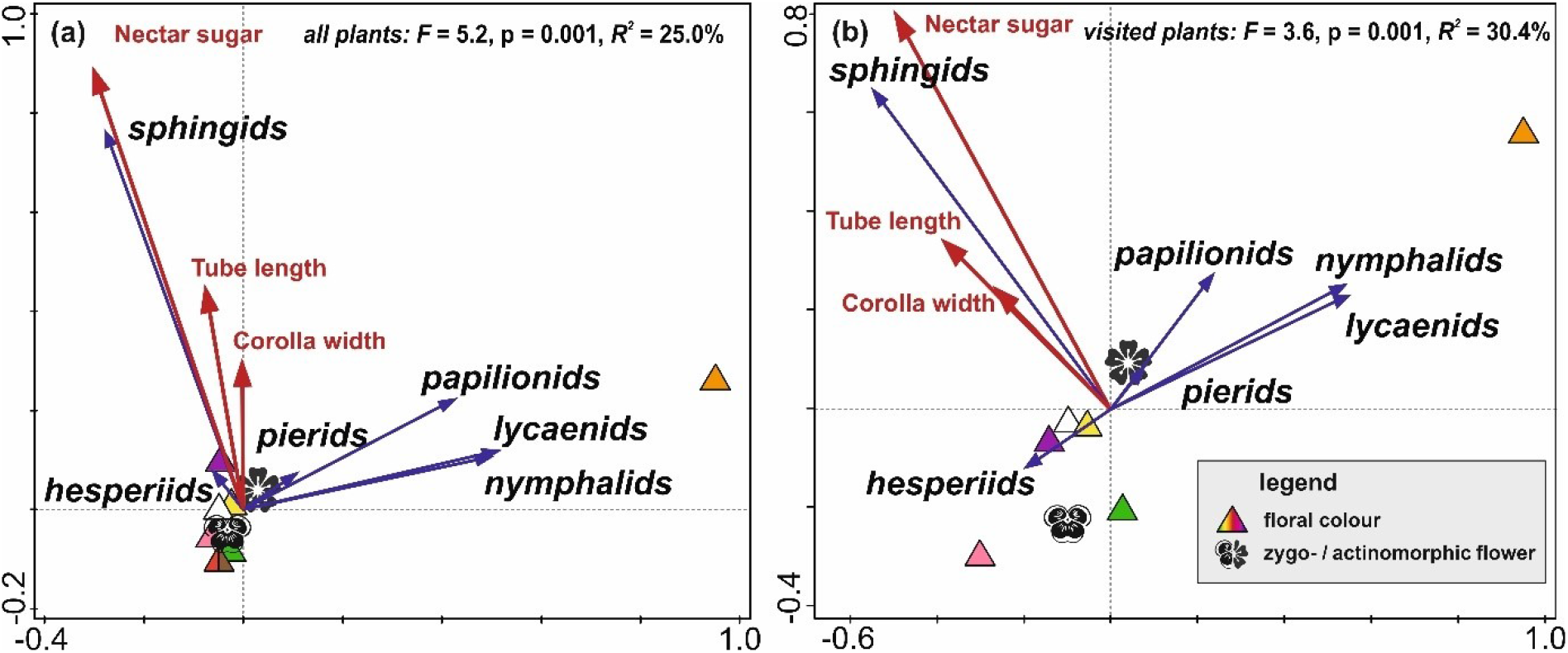
Redundancy analyses (RDA) revealing significant preferences of butterfly and sphingid families (represented by blue arrows) to floral traits (represented by red arrows and various symbols) on Mount Cameroon. The two RDA models were run for (a) all flowering plant species, and (b) the plant species visited by butterflies or moths.

## Discussion

### Importance of butterflies and sphingids as pollinators

On Mount Cameroon, butterflies and sphingids generally did not represent the most important pollinators as they collectively made up 4% of the flower-visiting community in tropical rainforests. Their numbers were dwarfed by flower-visiting bees, flies and beetles, representing 44.10%, 25.71% and 11.83% of visits of all recorded plants, respectively (Klomberg et al., 2020). The relative importance of lepidopterans in our uniquely comprehensive Afrotropical networks was even lower than in several partial networks from tropical forests of South-East Asia (e.g. Kato et al., 2008; Momose et al., 1998) and the Neotropics (e.g. Ramirez, 1989; Van Dulmen, 2001). In all these studies, flower visitation by lepidopterans was considerable (between 10-20% of all pollinators), although incomparable to bees (between 40-55%). We are not aware of any similar study from the Afrotropical forests.

Even though butterflies and sphingids visited about a third of all flowering plants in the study area, only a few plant species seemed to be primarily pollinated by these groups. Butterflies were the most common visitors of a single plant species, *Scadoxus cinnabarinus*, already known to be butterfly-pollinated (Mertens et al., 2020). In only a few other plant species, butterflies ranked high among all visiting groups (Table S4). Sphingids were not the most common visitors of any recorded plant species, but they ranked second visiting groups of *Anthocleista scandens*. Although plants within this genus have been reported as potentially pollinated by moths or sunbirds (Nsor & Chapman, 2013), its typically widely open chiropterophilous flowers (Weber et al., 2009) do not morphologically fit to sphingophily. However, several other plants commonly visited by butterflies (e.g. *Aframomum* spp. and *Cordia aurantiaca*) and sphingids (e.g. *Mussaenda tenuiflora* and *Clerodendrum silvanum*) in our study offer morphologically specialised flowers fitting to the lepidoptera-related pollination syndromes (Willmer, 2011). Their efficient pollination by lepidopterans can be expected from other studies of relative or similar plant species from other areas (e.g. Borges et al., 2003; Mizusawa et al., 2014). Several other plants commonly visited by the studied lepidopterans offered rather morphologically generalised inflorescences (e.g. *Distephanus biafrae*, *Melanthera scandens*, and *Crassocephalum montuosum*; all Asteraceae). Plants with such inflorescences were pollinated by butterflies or moths in some cases, although they were typically visited by rich pollinator communities and apparently did not only rely on lepidopterans (e.g. Budumajji & Solomon Raju, 2018; Valentin-Silva et al., 2016). Altogether, only a few plant species in our study seemed to depend on pollination by butterflies or sphingids based on the combination of their flowers’ morphology and visitation frequency. In conclusion, butterflies and sphingids seem to be relatively less relevant pollinators in tropical forests.

### Seasonal patterns of lepidopteran pollination

Pollination networks of butterflies and sphingids strongly differed between the studied seasons on Mount Cameroon. This was surely influenced by a plethora of factors affecting communities of both flower-visiting lepidopterans and flowering plants. The very high inter-seasonal turnover of both butterfly and moth species composition, as well as changes in species richness and abundance, were already reported in detail from Mount Cameroon (Maicher et al., 2018), as well as from other Afrotropical rainforests (e.g. Valtonen et al., 2013). Together with the confirmation of the high species turnover between dry and wet seasons (Fig. S2), this study also revealed seasonal changes in lepidopteran behaviour and networks of their pollination interactions. The general decline of flower-visiting butterflies is most probably connected to the local extreme precipitation during wet season. Adult butterflies and sphingids, as other large-winged insects, avoid the extreme rainfall on Mount Cameroon (Maicher et al., 2020). Simultaneously, the strong rains and related humidity also affect availability of nectar and its concentration (e.g. Aizen, 2003). The strong seasonality affects the flowering plant communities as well. Whereas many trees, usually offering large amounts of generally accessible nectar, flower in dry season, herbs and shrubs flowering in wet season are not able to replace this nectar production. Such unpublished suggestions were supported by the relatively high turnover of flowering plants between the two seasons (Fig. S2). Moreover plants flowering in the harsh conditions during wet season on Cameroon are often adapted to sunbirds (Bartoš & Janeček, 2014; Janeček et al., 2015). These could be both causes and consequences of the generally lower diversity and abundance of nectarivorous lepidopterans during wet season.

The prevailingly non-apparent trends in the plant-lepidopteran networks characteristics can be surprising because several other studies of pollination networks along elevation revealed diverse strong patterns (e.g. Classen et al., 2020; Ramos-Jiliberto et al., 2010). Nevertheless, we are not aware about any similar study on pollination by butterflies or sphingids along any tropical elevational gradient for comparison. Therefore, we hypothesise that lepidopteran species are surprisingly specialised for visited flowers, as visible in our relatively less connected and high specialised networks, and in the preferences for distinct floral traits. These preferences only partly change with environmental conditions (Klomberg et al., 2020). However, as discussed above, only a minor part of the visited flowers are specialised for lepidopteran pollination. Therefore, the significant trends in characteristics of the rather generalised plant-lepidopteran networks can hardly be expected. Nevertheless, only more studies of plant-lepidopteran networks from other tropical areas can challenge such hypotheses. We admit that for the only characteristic with a strong trend, nestedness, we do not have any apparent explanation, especially because the trends strongly differed inter-seasonally.

### Elevational patterns of lepidopteran pollination

Species richness of butterflies and sphingids showed the ‘low-elevation plateau with a mid-peak’ (*sensu* McCain & Grytnes, 2010), in accordance with numerous studies of tropical lepidopterans (e.g. Bärtschi et al., 2019; Maicher et al., 2020). The flower visitation frequencies of butterflies mirrored this result, whilst the importance of sphingids in pollination networks, both absolute and relative, increased along elevation. Because we are not aware of any studies on plant-lepidopteran pollination networks along any tropical elevational gradient for comparison, we can hardly speculate if the revealed patterns can be general. However, a study of the *Scadoxus cinnabarinus* pollination system showed peaking diversity and abundance of flower-visiting butterflies at mid-elevations of the plant range (Mertens et al., 2020). Besides numerous factors responsible for the generally high diversity at mid-elevations in many insect groups (e.g. Bärtschi et al., 2019; Beck et al., 2017), the patterns of plant-lepidoptera networks can be discussed in relation to the floral resources for nectarivorous butterflies and sphingids. Although we have no detailed data on the abundance of floral resources along the studied elevational gradient, the local diversity of trees linearly decreased along the gradient (Hořák et al., 2019). However, it remains questionable whether all flowering plants follow this pattern. The opposite trends of sphingid species richness and their importance in networks can be related to the dominance of a few highly mobile species among flower-visiting sphingids in all networks. Whilst their elevational diversity pattern was driven by numerous species with restricted elevational ranges on Mount Cameroon (Maicher et al., 2020), most of the identified sphingids in our networks were generalised long-distance vagrants (Table S1). Considering the generally small sizes of the plant-sphingid networks and the relatively smaller elevational variability, the revealed patterns could be caused by more or less random visitation by these mobile generalists.

### Traits in plant-lepidopterans networks

We found no evidence that longer proboscides or forewings of butterflies and sphingids elicited differences in the flower visitation behaviour, apart from the correlation between lengths of proboscis and floral tubes of flowers visited by hesperiids. The similar proboscis-tube relationship was already found in other studies (Bloch & Erhardt, 2008; Corbet, 2000; Johnson et al., 2017; Martins & Johnson, 2013), although such studies often found larger butterflies visiting larger flowers as well (e.g. Corbet, 2000; Tiple et al., 2009). Nevertheless, although preferring flowers with longer tubes, the long-proboscid lepidopterans were not more specialised in the meaning of their food-niche breadth (relative number of visited plant species) in our study. Therefore, we assume that even these morphologically specialised lepidopteran species were rather looking for any available resources in longer or deeper flowers which are more likely to be unreachable by other floral visitors, rather than being specialised for a few co-evolved plant species (cf. Martins & Johnson, 2013; Johnson et al., 2017).

Although we found no significant patterns among the morphological traits of lepidopterans along elevation when analysing all captured butterflies and sphingids, the lengths of forewing and proboscides showed U-shaped pattern when the analyses were restricted to the flower-visiting species. We expect that such patterns can be covered by other mechanisms when analysing complete lepidopteran community, including species with adults feeding on other resources than flowers. The increase of lepidopteran size towards the higher elevations was repeatedly reported and explained mostly by the need of larger area for basking in colder environments (e.g. Brehm et al. 2019). Nevertheless, the relatively larger bodies of lepidopterans in the lowest elevation seems surprising and difficult to explain.

Generally, preferences of butterflies and sphingids to the visited flowers were driven by floral colour, size, and nectar-sugar production in our study. This corroborates to numerous other studies (e.g. Tiple et al., 2009; Willmer, 2011). Moreover, the floral preferences neatly separated nocturnal sphingids from diurnal butterflies, as proposed by the pollination syndrome hypothesis (Willmer, 2011). Concurrent with other studies (Johnson et al., 2017; Kaczorowski et al., 2012), sphingids preferred longer and nectar-rich flowers on Mount Cameroon. Opposite to the sphingophilous syndrome (Willmer, 2011), they did not seem to prefer white flowers. Papilionids, nymphalids and pierids preferred orange flowers, the other floral traits were much less relevant. Strong preferences to floral colours were already shown for butterflies (e.g. Ômura & Honda, 2005; Pohl et al., 2011), and orange flowers are typical for the psychophilous plants (Willmer, 2011). Pierids and hesperiids expressed little to no preferences to any floral traits in this study. Such differences among butterfly families were already observed (Pohl et al., 2011; Yurtsever et al., 2010). Our detailed taxon-specific approach uncovered some of the limitations of the pollination syndrome hypothesis, especially that different traits can differ in their importance among particular syndromes, and that even individual subgroups of the single pollinator group can differ in their preferences (cf. Dellinger, 2020; Klomberg et al., 2020).

## Acknowledgments

We are grateful to Nestoral T. Fominka, Mercy Murkwe, Luma Francis Ewome, Raissa Dywou Kouede, Esembe Jacques Chi, Karolína Hrubá, Hernani Oliveira, Zuzana Sejfová, Pavel Kratochvíl, and other field assistants and students for help in the field; to Ivan Šonský, Sailee Sakhalkar, Petra Janečková, Eliška Chmelová, Marek Rybár, Jiří Hodeček, Jan Raška and several other people for help with processing the video-recordings; to Karolina and Ewelina Sroka for mounting of specimens; to Jan Wieringa and Naturalis staff for help with plant identification; to Eric B. Fokam and the Mount Cameroon National Park staff for their help with permits and logistics. This study was permitted by the Republic of Cameroon Ministries for Forestry and Wildlife and for Scientific Research and Innovation. Our research was funded by the Czech Science Foundation (16-11164Y) and Charles University (PRIMUS/17/SCI/8, UNCE204069 and GAUK 927116).

## Authors’ contributions

RT, ŠJ and JEJM conceived the ideas and designed methodology; JEJM, ŠJ, YK, INK and RT collected the plant-lepidopteran interactions data; JEJM, VM, SzS, RT, SD, PP, INK, LB and TP collected, identified and/or measured lepidopterans; ŠJ, YK and INK measured floral traits; JEJM and LB analysed data; JEJM and RT led the manuscript writing. All authors contributed to the drafts and approved for publication.

## Data Availability Statement

Data available via the Zenodo repository (*doi will be provided after acceptance*).

---

**Table S1.** List of all flower-visiting butterflies and sphingids identified from the video recordings on Mount Cameroon.

|  |  |  |
| --- | --- | --- |
| **Hesperiidae** | **Nymphalidae** | **Papilionidae** |
| *Acleros bibundica* | *Acraea alcinoe* | *Graphium polices* |
| *Andronymus magma* | *Acraea bonasia* | *Papilio charopus* |
| *Andronymus sp.1* | *Acraea elongate* | *Papilio cyproeofila* |
| *Apallaga alluaudi* | *Acraea epaea* | *Papilio dardanus* |
| *Apallaga intermixtus* | *Acraea judutta* | *Papilio hesperus* |
| *Apallaga meditrina* | *Acraea lycoa* | *Papilio menestheus* |
| *Apallaga mona* | *Acraea penelope* | *Papilio zenobia* |
| *Bettonula bettoni* | *Acraea pharsalus* | *Papilio zoroastres* |
| *Celaenorrhinus dargei* | *Acraea rogersi* |  |
| *Ceratrachia clara* | *Acraea tellus* | **Sphingids** |
| *Ceratrachia fako* | *Acraea umbra* | *Agrius convolvuli* |
| *Ceratrachia flava* | *Amauris damocles* | *Centroctena rutherfordi* |
| *Coeliades chalybe* | *Amauris echeria* | *Hippotion osiris* |
| *Coeliades forestan* | *Bebearia tentyrus* | *Hippotion sp.1* |
| *Coeliades hanno* | *Bicyclus anisops* | *Hippotion sp.2* |
| *Coeliades libeon* | *Bicyclus sciathis* | *Hippotion sp.3* |
| *Eagris decastigma* | *Cymothoe beckeri* | *Macroglossinae sp.1* |
| *Gorgyra sp.1* | *Cymothoe consanguis* | *Macroglossinae sp.2* |
| *Leona lissa* | *Cymothoe herminia* | *Macroglossinae sp.3* |
| *Meza sp.1* | *Cymothoe indamora* | *Macroglossinae sp.4* |
| *Osmodes lux* | *Cymothoe sangaris* | *Macroglossinae sp.5* |
| *Osmodes thora* | *Cymothoe weymeri* | *Macroglossinae sp.6* |
| *Paracleros sp.1* | *Euphaedra hewitsoni* | *Macroglossinae sp.7* |
| *Pardaleodes tibullus* | *Euphaedra losinga* | *Macroglossinae sp.8* |
| *Paronymus xanthias* | *Euphaedra sp.1* | *Macroglossum trochilus* |
| *Semalea sp.1* | *Euphaedra temerraria* | *Nephele accentifera* |
| *Tagiades flesus* | *Hypolimnas salmacis* | *Nephele sp.1* |
|  | *Junonia terea* | *Nephele sp.2* |
| **Lycaenidae** | *Phalanta eurytis* | *Nephele sp.3* |
| *Anthene definita* | *Precis milonia* | *Nephele sp.4* |
| *Cacyreus cf. lingens* | *Vanessula milca* | *Temnora iapygoides* |
| *Euchrysops malathana* |  | *Temnora sp.1* |
| *Hypolycaena sp.1* | **Pieridae** | *Theretra orpheus* |
| *Neurypexina lamprocles* | *Appias cf. sabina* | *Theretra sp.1* |
| *Thermoniphas alberici* | *Leptosia alcesta* | *Theretra sp.2* |
| *Zizula hylax* | *Leptosia cf. hybrida* | *Xanthopan morgani* |
|  | *Leptosia nupta* |  |

**Figure S1.**
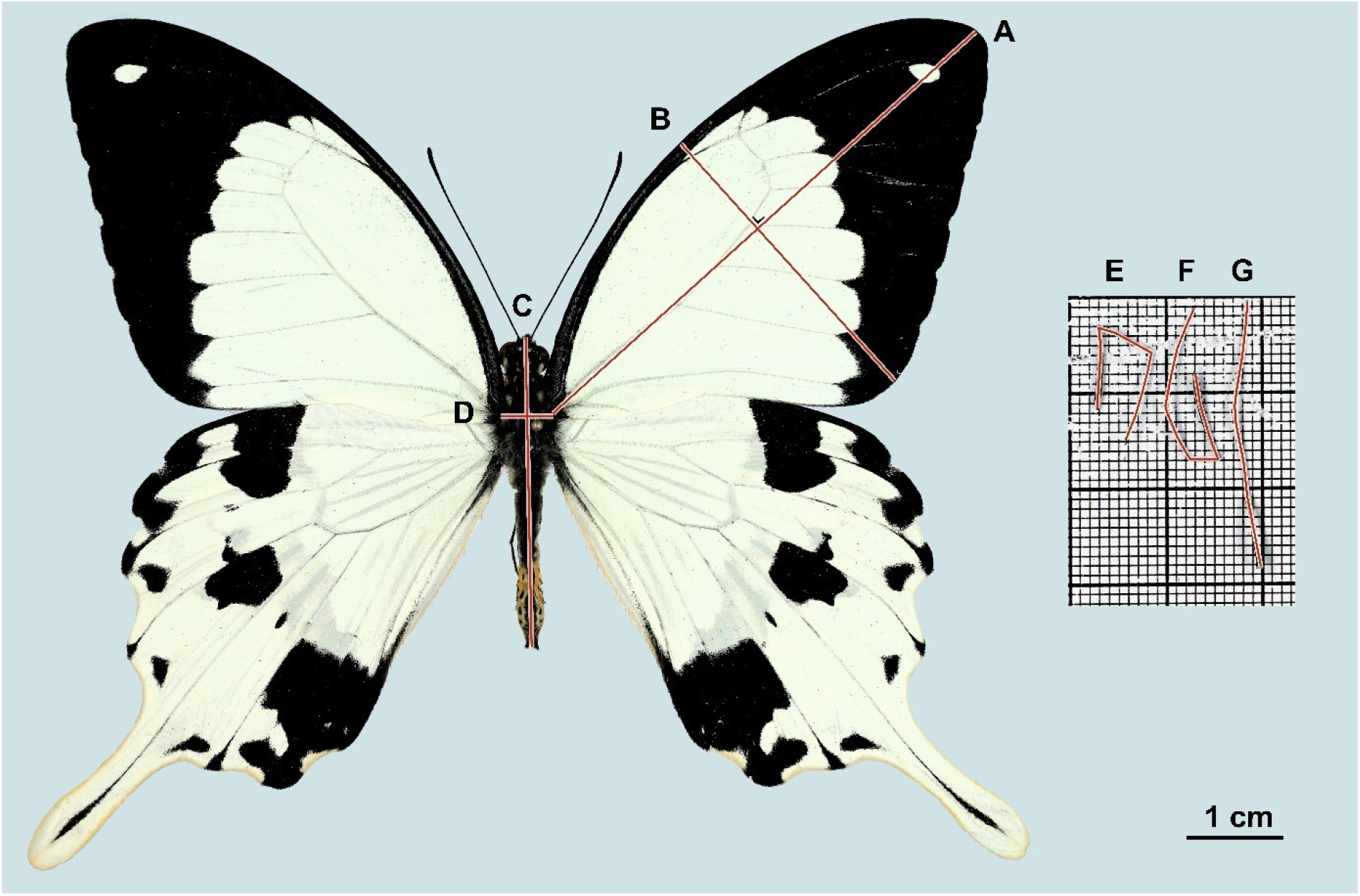
Illustration of the measured traits of flower-visiting butterflies and sphingids on Mount Cameroon. (A) Forewing length: from a wing base to a wingtip (defined as point where the tangent of a wing edge is perpendicular to the measure line). (B) Forewing width: perpendicular to the forewing length measure line and positioned so that the tangent of the outer wing margin is perpendicular to the measure line. (C) Body length: from the top of a head (excluding mouthparts) to the end of an abdomen (excluding genital valves). (D) Body width: measured where forewings attach to a thorax. (E-G) Lengths of fore-, mid-, and hindleg, respectively: from a base of tibia to the last tarsus (excluding tarsal claw), measured by a segmented line.

**Table S2.** Intercorrelation matrix of the measured traits of flower-visiting butterflies and sphingids on Mount Cameroon. After application of Bonferroni correction (39 analyses), only p < 0.001 were considered significant, these are highlighted by **bold**. All correlations were positive.

|  | Weight | Proboscis length | Forewing length | Forewing width | Body length | Body width | Foreleg length | Midleg length |
| --- | --- | --- | --- | --- | --- | --- | --- | --- |
| Proboscis length | $R^2 = 0.46$,<br>$p = 0.007$ | | | | | | | |
| Forewing length | $R^2 = 0.89$,<br>$p < 0.001$ | $R^2 = 0.42$,<br>$p = 0.014$ | | | | | | |
| Forewing width | $R^2 = 0.74$,<br>$p < 0.001$ | $R^2 = 0.36$,<br>$p = 0.037$ | $R^2 = 0.93$,<br>$p < 0.001$ | | | | | |
| Body length | $R^2 = 0.97$,<br>$p < 0.001$ | $R^2 = 0.56$,<br>$p = 0.001$ | $R^2 = 0.88$,<br>$p < 0.001$ | $R^2 = 0.74$,<br>$p < 0.001$ | | | | |
| Body width | $R^2 = 0.96$,<br>$p < 0.001$ | $R^2 = 0.49$,<br>$p = 0.004$ | $R^2 = 0.84$,<br>$p < 0.001$ | $R^2 = 0.68$,<br>$p < 0.001$ | $R^2 = 0.98$,<br>$p < 0.001$ | | | |
| Foreleg length | $R^2 = 0.89$,<br>$p < 0.001$ | $R^2 = 0.58$,<br>$p < 0.001$ | $R^2 = 0.94$,<br>$p < 0.001$ | $R^2 = 0.87$,<br>$p < 0.001$ | $R^2 = 0.92$,<br>$p < 0.001$ | $R^2 = 0.86$,<br>$p < 0.001$ | | |
| Midleg length | $R^2 = 0.90$,<br>$p < 0.001$ | $R^2 = 0.52$,<br>$p = 0.002$ | $R^2 = 0.94$,<br>$p < 0.001$ | $R^2 = 0.83$,<br>$p < 0.001$ | $R^2 = 0.92$,<br>$p < 0.001$ | $R^2 = 0.89$,<br>$p < 0.001$ | $R^2 = 0.92$,<br>$p < 0.001$ | |
| Hindleg length | $R^2 = 0.94$,<br>$p < 0.001$ | $R^2 = 0.41$,<br>$p = 0.019$ | $R^2 = 0.92$,<br>$p < 0.001$ | $R^2 = 0.82$,<br>$p < 0.001$ | $R^2 = 0.93$,<br>$p < 0.001$ | $R^2 = 0.88$,<br>$p < 0.001$ | $R^2 = 0.93$,<br>$p < 0.001$ | $R^2 = 0.88$,<br>$p < 0.001$ |

**Table S3.** Summary of butterflies and sphingids visiting of flowers / touching its reproductive organs at each sampling elevation and season on Mount Cameroon.

| Elevation<br>(m a.s.l.) | season | butterflies |  |  |  |  | sphingids | all<br>lepidopterans |
| --- | --- | --- | --- | --- | --- | --- | --- | --- |
|  |  | hesperiids | lycaenids | nymphalids | papilionids | pierids |  |  |
| 650 | dry | 69 / 65 | 3 / 3 | 22 / 21 | 19 / 19 | 12 / 12 | 13 / 13 | 138 / 133 |
| 1,100 | dry | 33 / 24 | 17 / 17 | 76 / 75 | 10 / 10 | 26 / 26 | 17 / 17 | 179 / 169 |
| 1,450 | dry | 11 / 2 | 10 / 10 | 11 / 11 | 7 / 7 | 141 / 137 | 29 / 28 | 209 / 195 |
| 2,200 | dry | / | 1 / 1 | / | 3 / 3 | 1 / 1 | 23 / 23 | 28 / 28 |
| 650 | wet | 60 / 57 | 13 / 13 | 10 / 10 | 1 / 1 | / | 10 / 10 | 94 / 91 |
| 1,100 | wet | 25 / 23 | / | 20 / 20 | 1 / 1 | / | 6 / 6 | 52 / 50 |
| 1,450 | wet | 4 / 4 | / | 20 / 20 | 1 / 1 | 1 / 1 | 4 / 4 | 30 / 30 |
| 2,200 | wet | / | / | / | / | / | 4 / 4 | 4 / 4 |
| Total |  | 628 / 595 |  |  |  |  | 106 / 105 | 734 / 700 |

**Table S4.** Plant species most visited by butterflies (top 10) and hawkmoths (top 5) on Mount Cameroon, as well as the rank of butterflies/sphingids among all functional groups of flower visitors (Klomberg et al. 2020).

| **butterflies** |  |  |  |
| --- | --- | --- | --- |
| **plant family** | **plant species** | **No. visits** | **rank** |
| Asteraceae | *Distephanus biafrae* | 136 | 2 |
| Zingiberaceae | *Aframomum* sp. 'purple' | 67 | 3 |
| Compositae | *Melanthera scandens* | 63 | 2 |
| Amaryllidaceae | *Scadoxus cinnabarinus* | 39 | 1 |
| Balsaminaceae | *Impatiens macroptera* | 27 | 3 |
| Vitaceae | *Cissus oreophylla* | 24 | 3 |
| Apocynaceae | *Tabernaemontana ventricosa* | 22 | 5 |
| Asteraceae | *Crassocephalum montuosum* | 22 | 5 |
| Boraginaceae | *Cordia aurantiaca* | 17 | 2 |
| Balsaminaceae | *Impatiens mannii* | 16 | 3 |
| **sphingids** |  |  |  |
| **plant family** | **plant species** | **No. visits** | **rank** |
| Gentianaceae | *Anthocleista scandens* | 15 | 2 |
| Asteraceae | *Distephanus biafrae* | 13 | 6 |
| Rubiaceae | *Mussaenda tenuiflora* | 11 | 3 |
| Apocynaceae | *Tabernaemontana ventricosa* | 8 | 7 |
| Lamiaceae | *Clerodendrum silvanum* | 7 | 4 |

**Figure S2.**
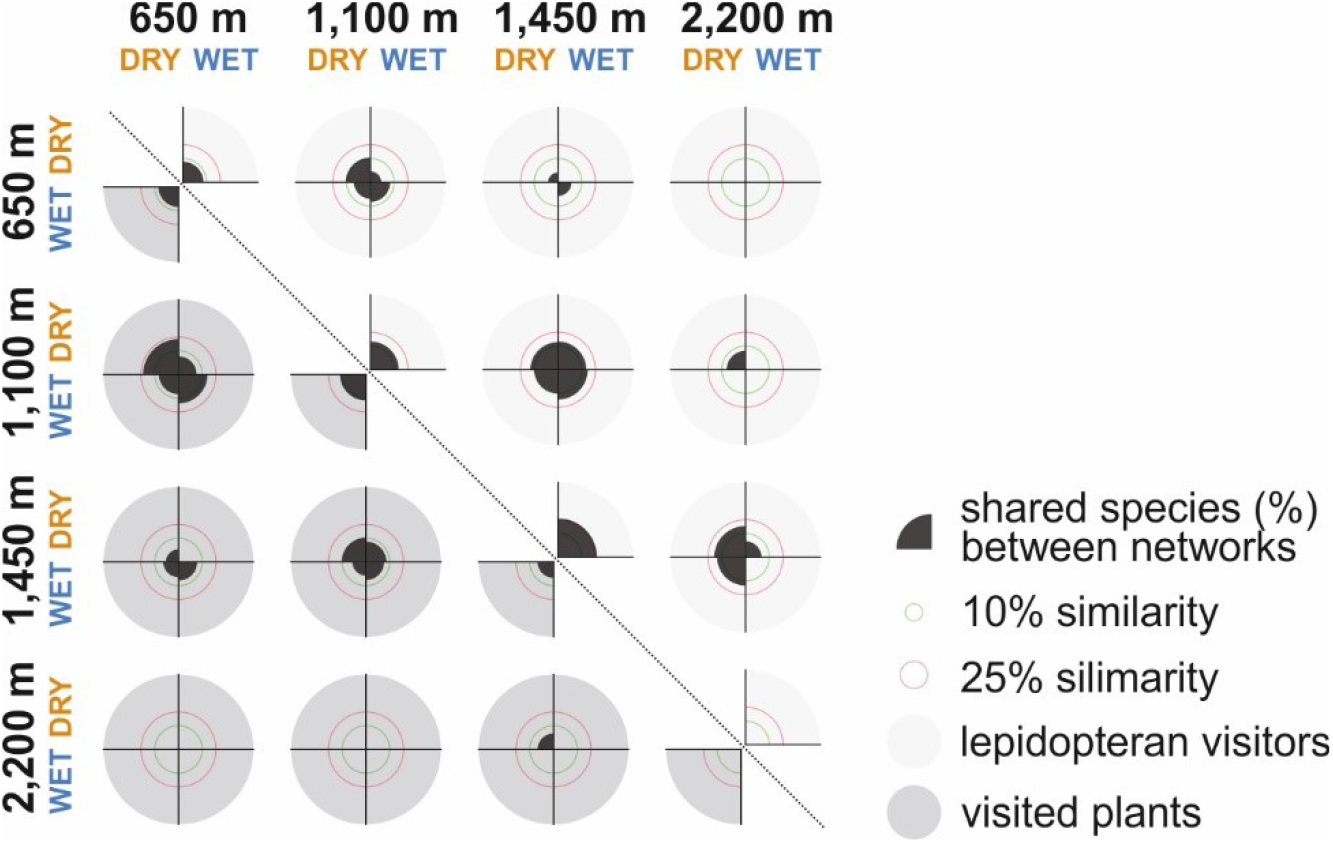
Turnover of lepidopteran and plant species in the studied plant-lepidoptera pollination networks on Mount Cameroon, visualised as proportions of the shared butterfly and sphingid visitor species and visited plant species among particular elevations and seasons.

**Table S5.**
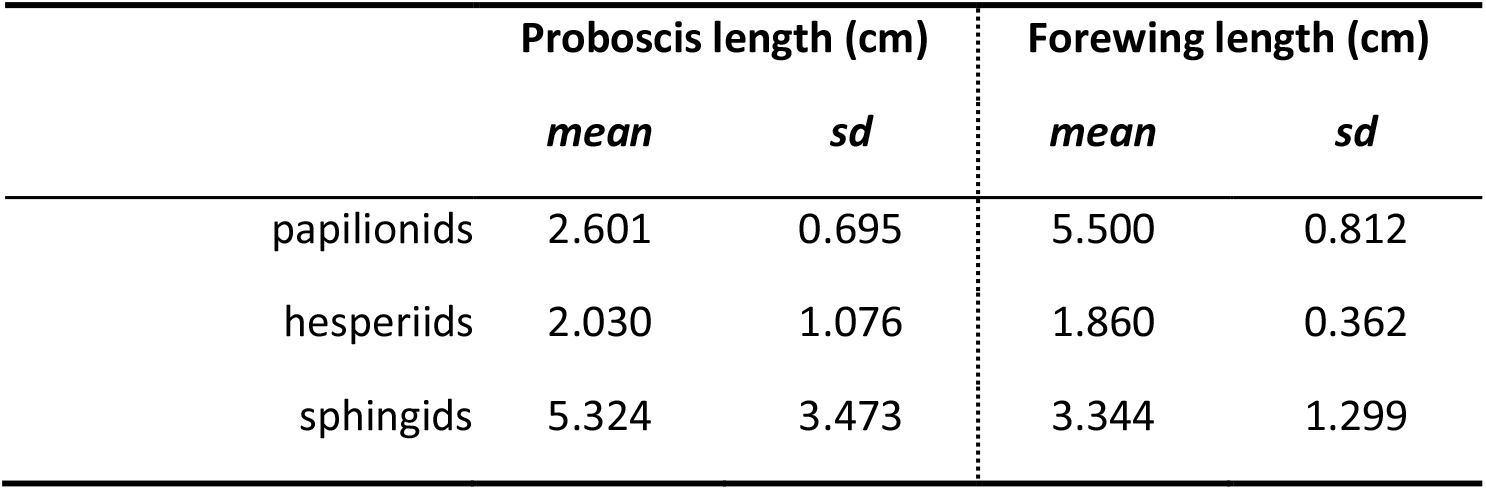
Summary of average trait values of butterflies and sphingids on Mount Cameroon.

|  | Proboscis length (cm) |  | Forewing length (cm) |  |
| --- | --- | --- | --- | --- |
|  | *mean* | *sd* | *mean* | *sd* |
| papilionids | 2.601 | 0.695 | 5.500 | 0.812 |
| hesperiids | 2.030 | 1.076 | 1.860 | 0.362 |
| sphingids | 5.324 | 3.473 | 3.344 | 1.299 |

**Table S6.** Results of linear models analysing effects of elevation on proboscis and forewing lengths of all butterflies and sphingids captured on Mount Cameroon.

|  | Proboscis length |  |  |  | Forewing length |  |  |  |
| --- | --- | --- | --- | --- | --- | --- | --- | --- |
|  | *df* | *F* | *p* | *R<sup>2</sup>adj* | *df* | *F* | *p* | *R<sup>2</sup>adj* |
| **all taxa** | 6 | 1.324 | 0.247 | 0.030 | 6 | 0.847 | 0.535 | 0.004 |
| **sphingids** | 6 | 0.626 | 0.709 | 0.052 | 6 | 1.130 | 0.356 | 0.088 |
| **hesperiids** | 6 | 0.720 | 0.634 | 0.033 | 6 | 0.925 | 0.480 | 0.039 |
| **papilionids** | 6 | 0.226 | 0.966 | 0.028 | 6 | 0.553 | 0.765 | 0.065 |

**Table S7.** Comparison of linear mixed-effect models (with family as random-effect variable) analysing effects of season, elevation, and their interaction on proboscis and forewing lengths of butterflies and sphingids visiting flowers on Mount Cameroon. Models with *ΔAICc* ≤ 2 from the most plausible model were considered as comparable

| | *residual*<br>*df* | *residual*<br>*deviance* | $\Delta AICc$ | *weight* | $R^2_{adj}$ |
| --- | --- | --- | --- | --- | --- |
| **Proboscis length** |  |  |  |  |  |
| season | 47 | 64.062 | 6.5 | 0.04 | 0 |
| **elevation** | **46** | **53.372** | **0** | **0.96** | **0.124** |
| season × elevation | 43 | 52.951 | 11.6 | 0 | 0.119 |
| **Forewing length** |  |  |  |  |  |
| season | 48 | 13.718 | 8.4 | 0.02 | 0 |
| **elevation** | **47** | **-0.486** | **0** | **0.98** | **0.046** |
| season × elevation | 44 | -4.651 | 11.7 | 0 | 0.119 |

